# “In-silico studies of facilitated VEGF(s) – VEGFR(s) bindings for assessment of Lysine as an indirect Low-Mol-Wt angiogen: Experimental validation of a potential synthetic Low-Mol-Wt angiogen”

**DOI:** 10.1101/077677

**Authors:** Priyanshu Verma, Aritri Bir, Anindita Banerjee, Joy Basu, Sujoy Kar, Sujoy Kumar Samanta, Debatosh Datta

## Abstract

In-vivo angiogenesis process is highly conserved and is mediated through a family of peptides having VEGF-A as the lead member. A respective receptor family comprising of members VEGFR-1, 2, 3 gets expressed on the endothelial cell membrane of the vascular bed in ischemic zone along with parallel expressions of VEGF-A, B, C, D and PlGF. Degree of ischaemia is the main regulator of these coupled expressions of angiogenic peptides/factors (AFs) and respective receptor(s) for a paracrine angiogenic process to take place. Physiological angiogenesis in intrauterine growth phase is the lead process in foetal growth, organogenesis and cellular specialization. Post birth and with aging, this process gets gradually inefficient and slow. In the present in-silico study, all angiogenic factors and receptor species are examined as for their binding stability in basal unaided condition and in presence of a possible Low-Mol-Wt linkage molecule–Lysine. Also a Lysine analogue 1,6-diaminohexanoic acid has been examined for its angiogenic potential both in dry docking experiment and in cell culture assay.

## 1. Introduction

VEGF is the major mitogen for endothelial cells. It has different isoforms. However, in case of human, VEGF family consists of mainly five isoforms - VEGF-A, VEGF-B, VEGF-C, VEGF-D and PlGF (placental growth factor) (Neufeld et al., 1999). All these isoforms have a specific role in the initiation of angiogenesis, which mainly exhibits the regeneration and growth of new blood vessels. In vascular exchange beds, angiogenesis, which is an extremely conserved process across species, forms basis of the supply line for healthy cells by providing them required nutrients and gas, signaling molecules and by removal of carbon dioxide and other metabolic end products (Ferrara and Davis-Smyth, 1997; Neufeld et al., 1999; Ferrara et al., 2003; Tammela et al., 2005; Olsson et al., 2006; Roskoski, 2007). Angiogenesis is endogenously/naturally controlled by a positive control feedback system where relative hypoxia/ischaemia of an organ or tissue acts as a positive modifier of inducible angiogenic process through elaboration of hypoxia induced factors and VEGF family of proteins. This inducible system is considerably complex and precise, which involve both endogenous inhibitors and activators (Roskoski, 2007). Inducible angiogenic process is an absolute necessity and supports intrauterine growth, including organogenesis, rapid cellular division and differentiation. However, with aging “normal physiological angiogenic process” gets more and more inadequate for required cellular growth, replenishments and continuous repair processes. Occasional very high demand of the process for tissue repair and regeneration also goes mostly unattended. In-vivo angiogenesis is promoted through enhanced production of VEGF(s) by endothelial cells in relative ischaemic tissues followed by its binding to the dedicated VEGFR(s) in the immediate vicinity in a paracrine effect. In addition, the VEGFR family consists of two non-protein kinases (Neuropilin-1 and 2) and three receptor tyrosine kinases (VEGFR-1, VEGFR-2 and VEGFR-3) (Neufeld et al., 1999; Tammela et al., 2005; Olsson et al., 2006; Roskoski, 2007). VEGFR-1 binds to VEGF-A, VEGF-B and PlGF; VEGFR-2 binds to VEGF-A, VEGF-C and VEGF-D; VEGFR-3 binds to VEGF-C and VEGF-D. VEGFR-1 and VEGFR-2 receptors are mainly found on the blood vascular endothelium and responsible for angiogenesis. However, VEGFR-3 receptors are mainly found on the lymphatic endothelium and responsible for lymphangiogenesis (Ferrara and Davis-Smyth, 1997; Ferrara et al., 2003; Tammela et al., 2005; Kasap and Sazci, 2008) (See Fig. 1).

**Fig. 1.**
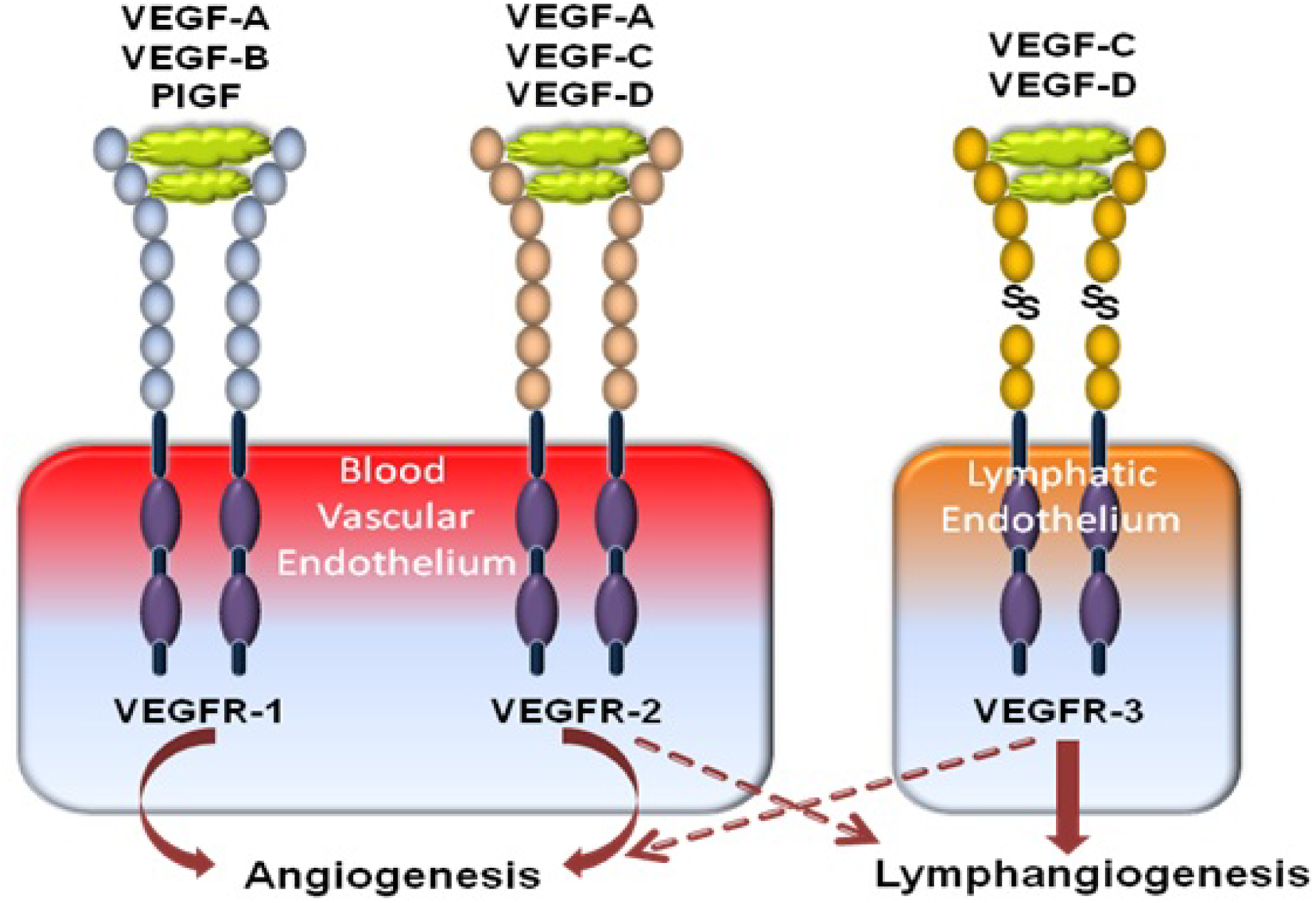
Schematic representation of the VEGF family receptors and VEGF isoform(s) interaction.

Since uncontrolled non-hierarchical tumor angiogenesis plays central role(s) in proliferation of cancerous cells, huge majority of the studies are mainly focused on possible inhibition processes of it (Luo et al., 2009; Ferrara and Adamis, 2016; Kareva, 2016; Soto-Ortiz and Finley, 2016).

On the contrary, the area of inducible augmented controlled angiogenesis for reperfusion of ischaemic tissues is substantially slow and almost a non-starter since the clinical limitations of VEGF(s) as therapeutic agent(s) became a reality (Deveza et al., 2012; Moserle et al., 2014).

As mentioned clinical limitation of VEGFs as a therapeutic agent family has made the area of controlled angiogenesis a slowly evolving one, because of the absence of a candidate molecule with a good safety profile and reproducible angiogenic potential compared to rapid progresses made in suppression of it (Ferrara and Adamis, 2016).

In this study binding stability of different VEGFs and VEGFR(s) have been examined both in presence and absence of the putative molecular binder –Lysine, with the help of in-silico docking tools. In addition, the potential of one carbon elongated (one additional −CH_2_ group in the carbon chain) analogue Lysine molecule as a potent angiogen has been examined through in-vitro cell culture experiments followed by in-silico validation.

## 2. Materials and Methods

### 2.1. In-silico docking analysis of different Lysine-VEGFs-VEGFRs combinations

This section is mainly focused on the in-silico analysis of binding affinity of different VEGFs to VEGFR(s) in presence of basic amino acid Lysine (Datta et al., 2001). Docking tools -Z-DOCK (Pierce et al., 2014) and PATCHDOCK (Schneidman-Duhovny et al., 2005) followed by FIREDOCK (Andrusier et al., 2007; Mashiach et al., 2008) has been used for the in-silico studies. Required protein molecules’ pdb files have been selected from the PDB database (Berman et al., 2000) and further modified for the removal of pre-existing ligands and other foreign compounds using Chimera tool. Table 1 represents the molecules with their PDB IDs that have been used for the in-silico docking experiments. The Lysine molecules’ sdf file has been downloaded from the PubChem server (Kim et al., 2016) and further converted to pdb extension using an offline tool-Open Babel (O’Boyle et al., 2011).

**Table 1.** Molecules with their PDB IDs that have been used in the present study.

| S. No. | Family | Name | PDB ID | Peptide Length | Modification Required* |
| --- | --- | --- | --- | --- | --- |
| 1 | VEGFRs | VEGFR-1 | 3HNG | 360 aa | Yes, 2 ligands removed |
| 2 |  | VEGFR-2 | 1Y6A | 366 aa | Yes, 2 ligands removed |
| 3 |  | VEGFR-3 | 4BSJ | 232 aa | Yes, 1 ligand removed |
| 4 | VEGFs | VEGF-A | 1VPF | 102 aa | No, used as it is |
| 5 |  | VEGF-B | 2C7W | 99 aa | Yes, 1 ligand removed |
| 6 |  | VEGF-C | 4BSK | 121 aa | Yes, 1 ligand + VEGFR-3 removed |
| 7 |  | VEGF-D | 2XV7 | 112 aa | Yes, 3 ligands removed |
| 8 |  | PIGF | 1FZV | 132 aa | Yes, 1 ligand removed |
| *Peptide modification required before proceeding for docking studies |  |  |  |  |  |

Prediction studies have been performed for the finding of most probable Lysine binding site(s) on VEGFs and VEGFR(s) using Z-DOCK online tool. However, for stability analyses, only FIREDOCK results have been considered. Chimera 1.10.2 (Pettersen et al., 2004) and Swiss-PdbViewer 4.1.0 (Guex and Peitsch, 1997) have been used for visualization and editing of models generated through docking studies.

Present study is mainly focused on a hypothesis – basic amino acid Lysine or elongated analogue(s) of the molecule (upto a certain chain length which needs to be ascertained) possibly acts as a molecular binder between the in-vivo angiogenic VEGF peptide(s) and their receptor(s) (VEGFRs) thereby augmenting binding stability between the two, resulting in induction of enhanced biological response – angiogenesis (See Fig. 2). This facilitated binding resulting in enhanced induced angiogenesis help(s) in controlled reperfusion of ischaemic tissues.

**Fig. 2.**
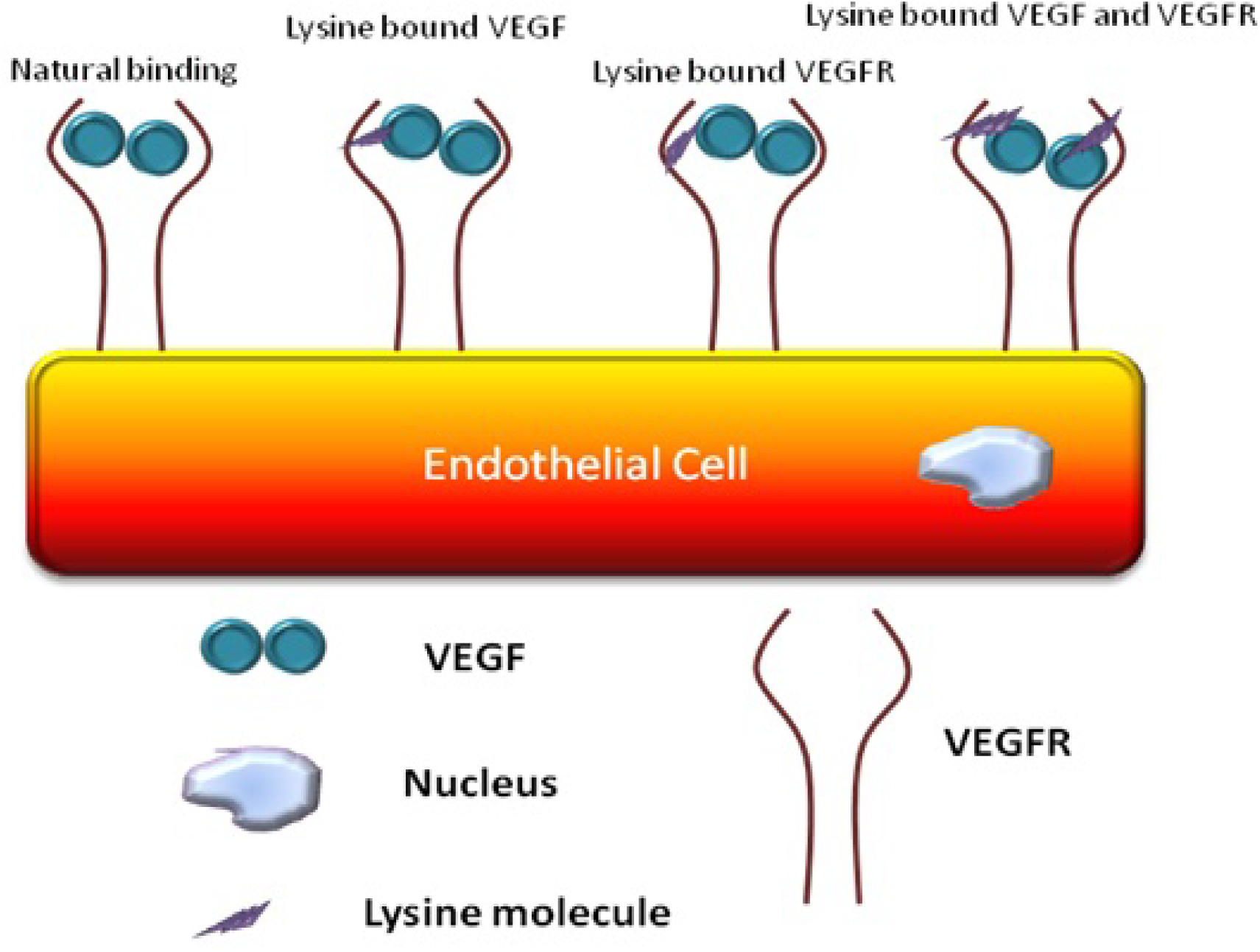
Hypothetical schematic representation of binding of different Lysine–VEGF-VEGFR combinations on endothelial cell.

Present study design includes different modes of approach to validate the hypothesis using in-silico binding studies:

i. VEGFR-X (X= 1, 2, 3) are presented for binding with VEGF-Y (Y=A, B, C, D) and PlGF as per their binding affinities (as shown in Fig. 1 and Table 2) and considered as natural or reference step.
ii. Lysine being coupled to VEGFR-X (X= 1, 2, 3) to generate the L–VEGFR-X complex, respectively.
iii. These complexes are then presented for binding with VEGF-Y (Y=A, B, C, D) and PlGF as per their binding affinities (as shown in Fig. 1 and Table 2).
iv. Lysine is first coupled to VEGF-Y (Y=A, B, C, D) and PlGF to generate the L– VEGF-Y and L–PlGF complexes, respectively. Similarly, these complex peptides are then presented for binding studies with VEGFR-X (X= 1, 2, 3) as per their binding affinities (as shown in Fig. 1 and Table 2).
v. All Lysine coupled L–VEGF-Y (Y=A, B, C, D) and L–PlGF complexes are presented to Lysine coupled L–VEGFR-X (X= 1, 2, 3) complex as per their binding affinities (as shown in Fig. 1 and Table 2).

**Table 2.** Different combinations used for docking studies of VEGFs and VEGFRs as per their binding affinities.

|  | VEGFR-1 | VEGFR-2 | VEGFR-3 |
| --- | --- | --- | --- |
| VEGF-A | (i) VEGFR-1 + VEGF-A | (i) VEGFR-2 + VEGF-A |  |
|  | (ii) L-VEGFR-1 + VEGF-A | (ii) L-VEGFR-2 + VEGF-A |  |
|  | (iii) VEGFR-1 + L-VEGF-A | (iii) VEGFR-2 + L-VEGF-A |  |
|  | (iv) L-VEGFR-1 + L-VEGF-A | (iv) L-VEGFR-2 + L-VEGF-A |  |
| VEGF-B | (i) VEGFR-1 + VEGF-B |  |  |
|  | (ii) L-VEGFR-1 + VEGF-B |  |  |
|  | (iii) VEGFR-1 + L-VEGF-B |  |  |
|  | (iv) L-VEGFR-1 + L-VEGF-B |  |  |
| VEGF-C |  | (i) VEGFR-2 + VEGF-C | (i) VEGFR-3 + VEGF-C |
|  |  | (ii) L-VEGFR-2 + VEGF-C | (ii) L-VEGFR-3 + VEGF-C |
|  |  | (iii) VEGFR-2 + L-VEGF-C | (iii) VEGFR-3 + L-VEGF-C |
|  |  | (iv) L-VEGFR-2 + L-VEGF-C | (iv) L-VEGFR-3 + L-VEGF-C |
| VEGF-DDD |  | (i) VEGFR-2 + VEGF-D | (i) VEGFR-3 + VEGF-D |
|  |  | (ii) L-VEGFR-2 + VEGF-D | (ii) L-VEGFR-3 + VEGF-D |
|  |  | (iii) VEGFR-2 + L-VEGF-D | (iii) VEGFR-3 + L-VEGF-D |
|  |  | (iv) L-VEGFR-2 + L-VEGF-D | (iv) L-VEGFR-3 + L-VEGF-D |
| PIGF | (i) VEGFR-1 + PIGF |  |  |
|  | (ii) L-VEGFR-1 + PIGF |  |  |
|  | (iii) VEGFR-1 + L-PIGF |  |  |
|  | (iv) L-VEGFR-1 + L-PIGF |  |  |
|  | Natural process (Yellow) | Lysine combined complex (Green) | Affinity not found in nature (Red) |

### 2.2. Angiogenic potential of elongated Lysine molecule (ELM) using cell culture assay

The Angiogenic potential of the ELM was tested experimentally by binding of human VEGF to its putative receptors. Human malignant lung epithelial cells (A-549) were grown on cover-slip in DMEM/F-12 (1:1) or RPMI 1640 media supplemented with 10% FBS (Fetal Bovine Serum), insulin (0.1 units/ml), L-glutamine (2 mM), sodium pyruvate (100 g/ml), non-essential amino acids (100 μM), penicillin (100 units/ml) and streptomycin (100 μg/μl). Cells were incubated at 37 °C in a humidified atmosphere of 5% CO_2_. The pH of the medium was maintained at 6.2 for the binding assay of VEGF to its receptor in the presence of ELM. Cells were pre-treated with media alone or with 25 μg/ml of ELM for 30 min. Cells were then incubated with 10 ng/ml human VEGF for further 30 min at 37 °C in a humidified atmosphere of 5% CO_2_. Cells were fixed and stained with anti-VEGF antibody coupled with Alexaflour 488 and were visualized under Zeiss confocal microscope.

In-silico docking study for ELM (elongated by one –CH_2_ residue only) had also been performed using the binding affinity of VEGF-A and VEGFR-1 as a reference.

## 3. Results and Discussions

### 3.1. In-silico docking analysis of different VEGF-VEGFR combinations in presence and in absence of Lysine

Z-DOCK results for all VEGFs and VEGFRs show differentially distributed preferential binding sites for the external Lysine molecule (Fig. 3). While VEGF-A and VEGF-B have well demarcated binding sites for external Lysine molecule, VEGF-C, VEGF-D and PlGF show rather distributed locations of preferred binding. Similarly VEGFR-1 shows rather confined preferred locations of binding rather than VEGFR-2 and VEGFR-3.

**Fig 3.**
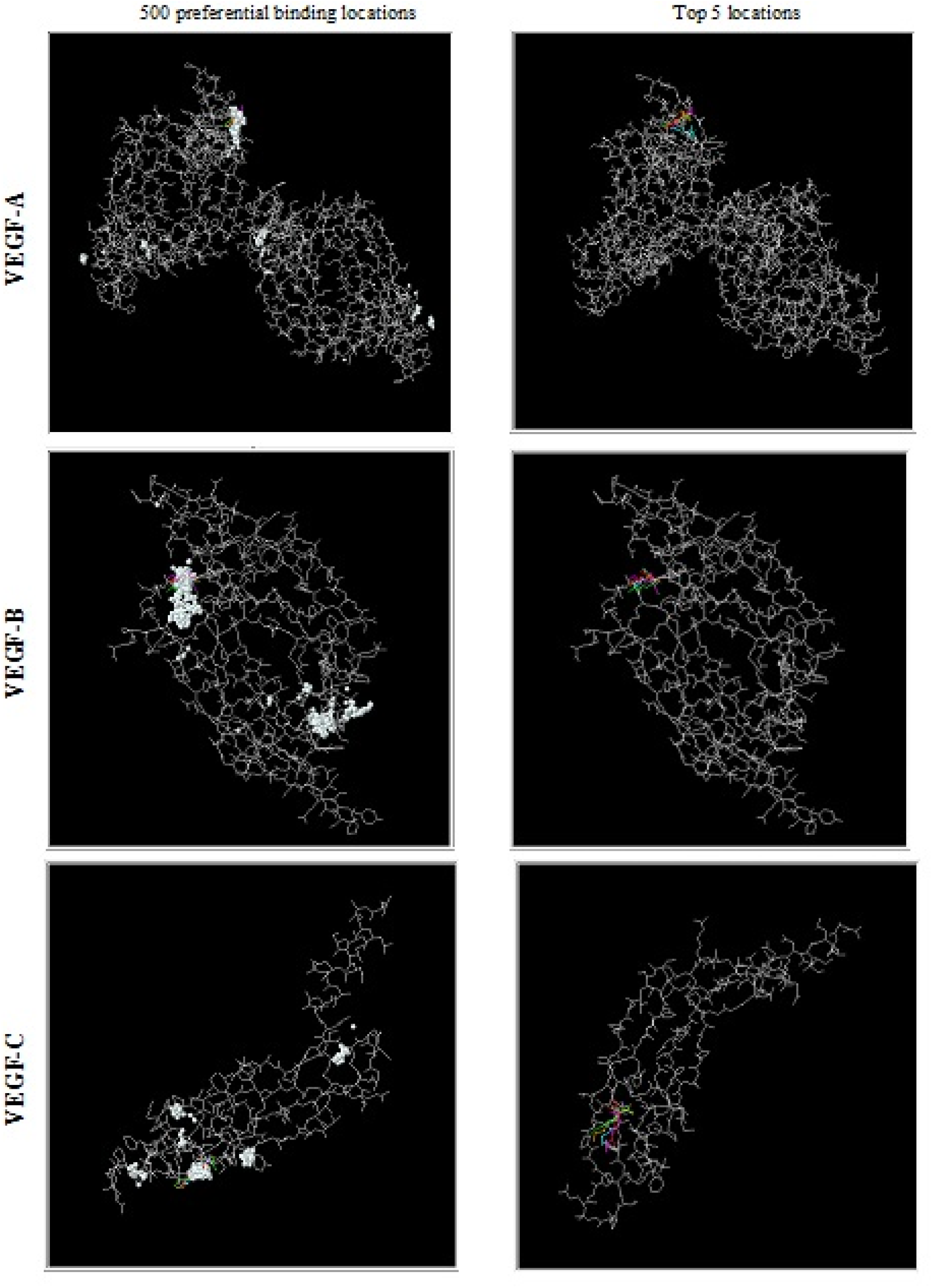
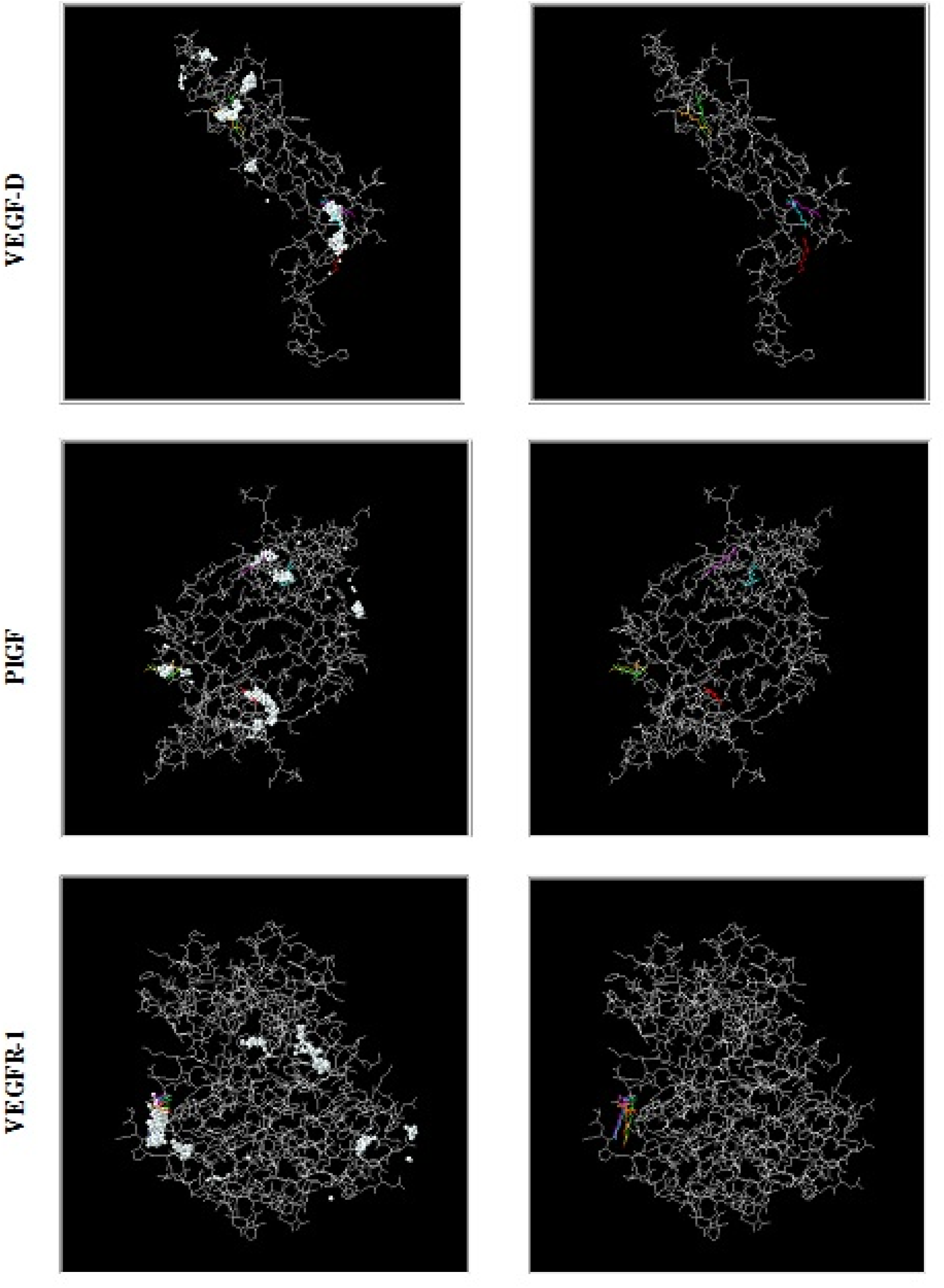
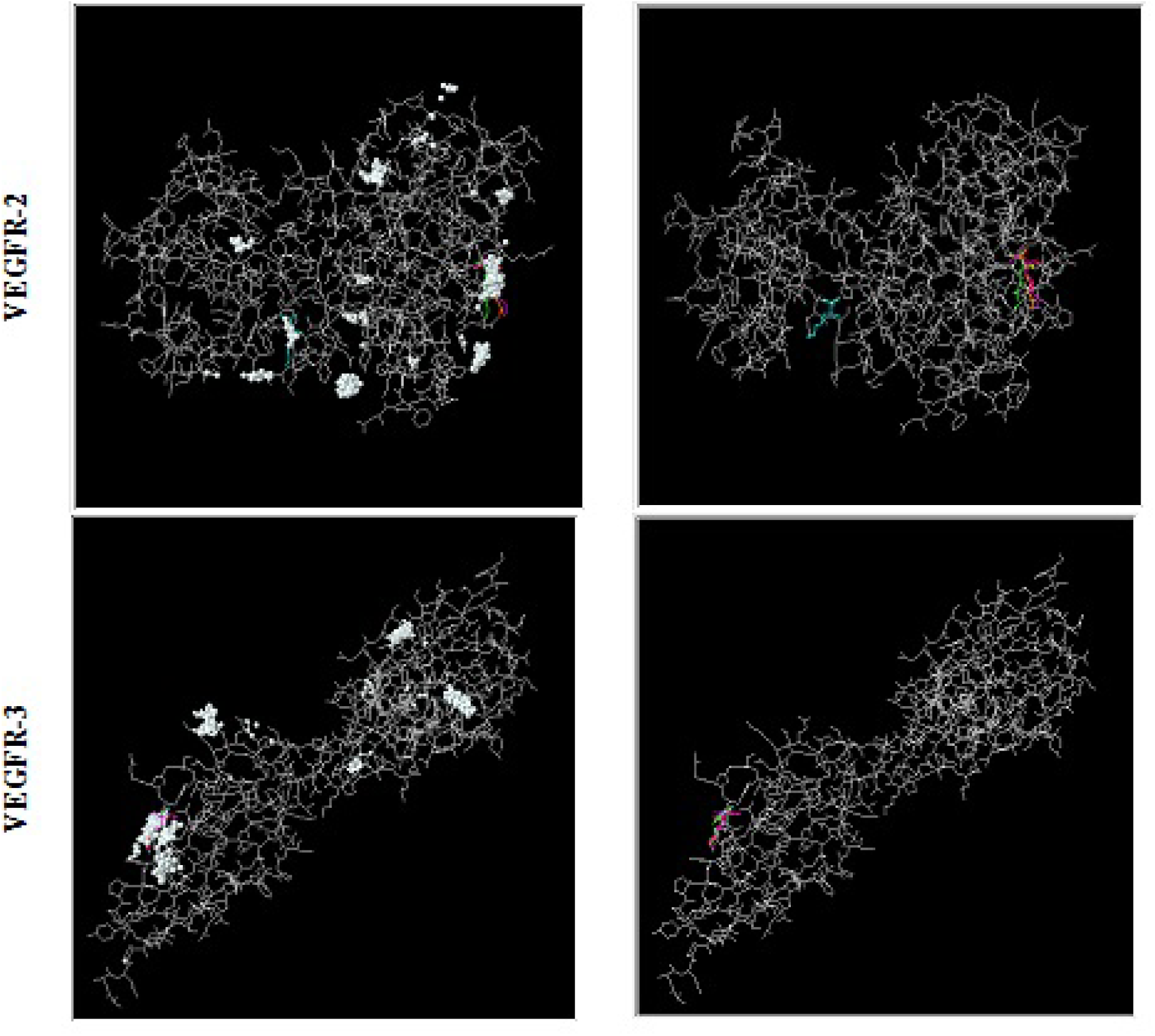
Z-DOCK results showing top 500 preferential binding locations (white spheres) and top 5 locations (colored lines) for externally added Lysine molecule as a ligand in all VEGFs and VEGFRs.

While these locations of preferential binding of externally added lysine molecule do raise questions and possibilities as to the rationality of very existence of such locations at all, added lysine did show very remarkable augmentation of binding between the two peptides (unqualified VEGF and VEGFR), in-vitro (Fig. 4) possibly signifying a thus-far unexplained relationship between the added molecular species and the two-peptides’ binding dynamics.

**Table 3.**
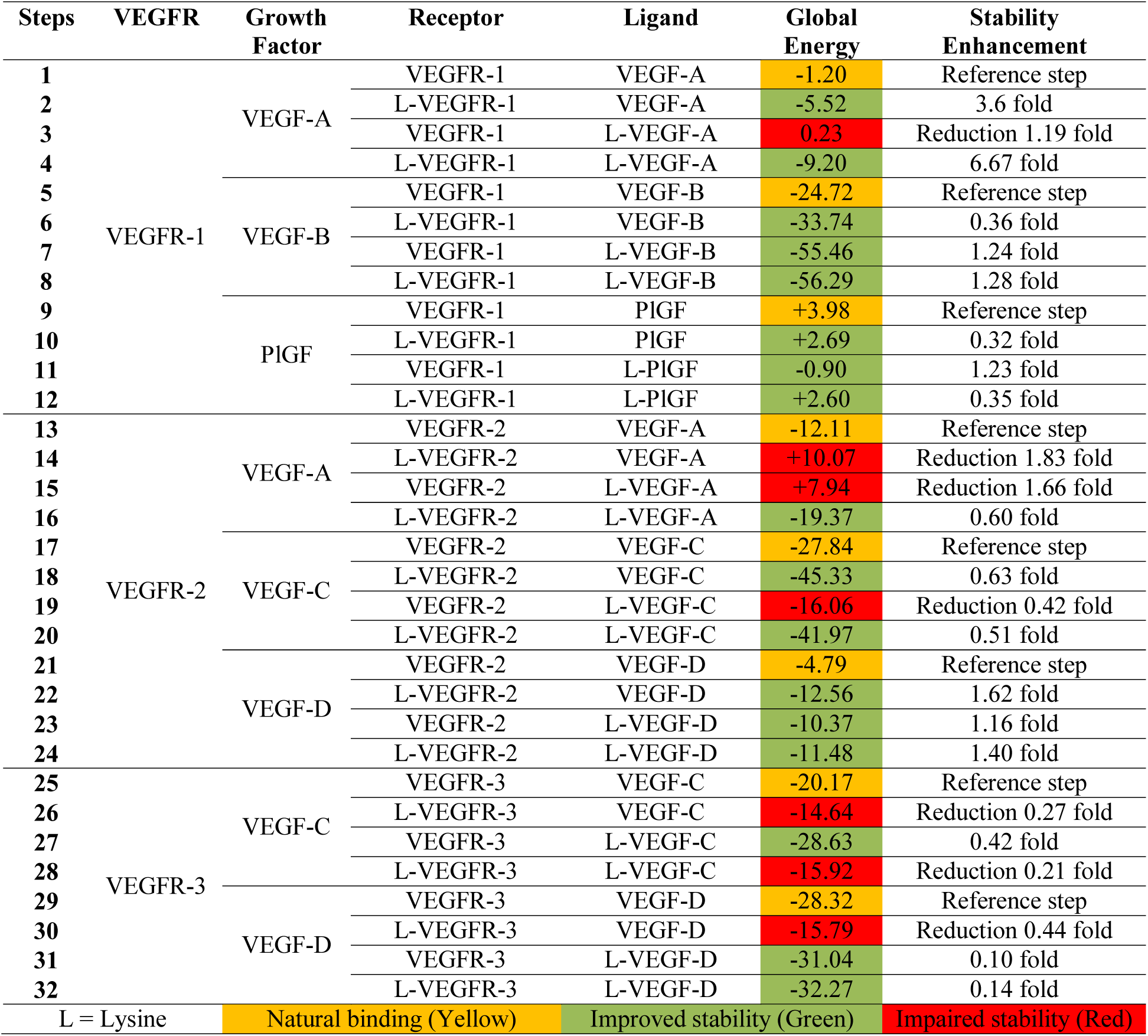
Overall docking studies used for different possible VEGFs–VEGFRs combinations.

| Steps | VEGFR | Growth Factor | Receptor | Ligand | Global Energy | Stability Enhancement |
| --- | --- | --- | --- | --- | --- | --- |
| 1 | VEGFR-1 | VEGF-A | VEGFR-1 | VEGF-A | -1.20 | Reference step |
| 2 |  |  | L-VEGFR-1 | VEGF-A | -5.52 | 3.6 fold |
| 3 |  |  | VEGFR-1 | L-VEGF-A | 0.23 | Reduction 1.19 fold |
| 4 |  |  | L-VEGFR-1 | L-VEGF-A | -9.20 | 6.67 fold |
| 5 |  | VEGF-B | VEGFR-1 | VEGF-B | -24.72 | Reference step |
| 6 |  |  | L-VEGFR-1 | VEGF-B | -33.74 | 0.36 fold |
| 7 |  |  | VEGFR-1 | L-VEGF-B | -55.46 | 1.24 fold |
| 8 |  |  | L-VEGFR-1 | L-VEGF-B | -56.29 | 1.28 fold |
| 9 |  | PIGF | VEGFR-1 | PIGF | +3.98 | Reference step |
| 10 |  |  | L-VEGFR-1 | PIGF | +2.69 | 0.32 fold |
| 11 |  |  | VEGFR-1 | L-PIGF | -0.90 | 1.23 fold |
| 12 |  |  | L-VEGFR-1 | L-PIGF | +2.60 | 0.35 fold |
| 13 | VEGFR-2 | VEGF-A | VEGFR-2 | VEGF-A | -12.11 | Reference step |
| 14 |  |  | L-VEGFR-2 | VEGF-A | +10.07 | Reduction 1.83 fold |
| 15 |  |  | VEGFR-2 | L-VEGF-A | +7.94 | Reduction 1.66 fold |
| 16 |  |  | L-VEGFR-2 | L-VEGF-A | -19.37 | 0.60 fold |
| 17 |  | VEGF-C | VEGFR-2 | VEGF-C | -27.84 | Reference step |
| 18 |  |  | L-VEGFR-2 | VEGF-C | -45.33 | 0.63 fold |
| 19 |  |  | VEGFR-2 | L-VEGF-C | -16.06 | Reduction 0.42 fold |
| 20 |  |  | L-VEGFR-2 | L-VEGF-C | -41.97 | 0.51 fold |
| 21 |  | VEGF-D | VEGFR-2 | VEGF-D | -4.79 | Reference step |
| 22 |  |  | L-VEGFR-2 | VEGF-D | -12.56 | 1.62 fold |
| 23 |  |  | VEGFR-2 | L-VEGF-D | -10.37 | 1.16 fold |
| 24 |  |  | L-VEGFR-2 | L-VEGF-D | -11.48 | 1.40 fold |
| 25 | VEGFR-3 | VEGF-C | VEGFR-3 | VEGF-C | -20.17 | Reference step |
| 26 |  |  | L-VEGFR-3 | VEGF-C | -14.64 | Reduction 0.27 fold |
| 27 |  |  | VEGFR-3 | L-VEGF-C | -28.63 | 0.42 fold |
| 28 |  |  | L-VEGFR-3 | L-VEGF-C | -15.92 | Reduction 0.21 fold |
| 29 |  | VEGF-D | VEGFR-3 | VEGF-D | -28.32 | Reference step |
| 30 |  |  | L-VEGFR-3 | VEGF-D | -15.79 | Reduction 0.44 fold |
| 31 |  |  | VEGFR-3 | L-VEGF-D | -31.04 | 0.10 fold |
| 32 |  |  | L-VEGFR-3 | L-VEGF-D | -32.27 | 0.14 fold |
| L = Lysine |  | Natural binding (Yellow) |  | Improved stability (Green) |  | Impaired stability (Red) |

Table 3 indicates overall docking profiles of different possible VEGF –VEGFR combinations (Table 2 and Fig. 1) that have been used in the present study and which all influence angiogenesis as well as lymphangiogenesis processes. Complex having lowest global energy among the docked results/structures has been considered as the best solution for respective docking profile (Table 3).

#### 3.1.1. VEGFR-1 – VEGF-A vs. VEGF-B binding

Binding stability of VEGF-A and VEGF-B with VEGFR-1 shows a very interesting direction. While VEGF-A binding to VEGFR-1 is substantially less stable than VEGF-B to VEGFR-1 (going by global energy consideration of −1.20 Units vs −24.72 Units), added lysine (1:1) to VEGF-A, VEGF-B and VEGFR-1 modified the stability remarkably. While VEGF-A➔VEGFR-1 complex showed an enhancement of stability by ~650%, VEGF-B➔ VEGFR-1 complex was more stable by about 120% (Table 3).

In-vivo physiological roles of VEGF-A and VEGF-B in induction of “normal” angiogenic process is more biased towards VEGF-A although VEGF-B shows much more stability in unaided binding, which should have resulted in enhanced parallel physiological response compared to VEGF-A (Bry et al., 2014; Zhang et al., 2014). This differential binding stability in favour of VEGF-B but not resulting in equivalent biological response remains unexplained.

#### 3.1.2. PlGF – VEGFR-1 binding

VEGFR-1–PlGF complex seems physiologically insignificant as for angiogenesis process is concerned, even though the global energy profile was reduced in presence of Lysine attached to both PlGF and VEGFR-1. Assigned role(s) of PlGF in establishment of placental circulation and maintenance throughout fetal growth remains to be quantified (Kim et al., 2012). However, fetal growth issues like Small-for-Date babies, IUGR, Spontaneous Abortions without Uterine structural abnormalities or hormonal aberrations need be probed through this apparently new direction.

#### 3.1.3. VEGFR-2 binding

Unaided VEGFR-2–VEGF-C binding tops the three components -VEGF-A, C and D (in terms of stability) with VEGF-B binding to VEGFR-1 running very close. Unaided initial binding in VEGFR-2–VEGF-C complex is much more stable than the other two unaided (physiological) bindings, thereby possibly indicating a much more pronounced angiogenic role of VEGFR-2– VEGF-C complex, nearly parallel to VEGFR-1–VEGF-B complex (Chen et al., 2013).

VEGFR-2–VEGF-D demonstrates nearly 140 % more stable complexes in the presence of Lysine in both receptor and ligands, compared to unaided baseline physiological binding profiles. In cases of VEGFR-2 bindings to VEGF-A and VEGF-C enhancements varied between 50 – 60 %. Any exploitable clinical outcome and significance of this enhanced stability of VEGF-D – VEGFR-2 complex in the presence of an added amino acid molecule remains to be examined.

#### 3.1.4. VEGFR-3 binding

Binding augmentation and final complex stability enhancements in VEGFR-3–VEGF-C and VEGFR-3–VEGF-D are not comparable to VEGFR-2–VEGF-C, A or D complexes. Significance of presence of VEGFR-3 may possibly lie in ontogeny of receptor series without much of physiological role(s). It would be interesting to note the development/appearance sequence of different receptor species in terms of time. Physiologically, these bindings may not signify much in terms of induced angiogenic responses in controlled reperfusion of ischemic tissues (even in the presence of the known augmentor Lysine).

### 3.2. Angiogenic potential of elongated Lysine molecule (ELM) using cell culture assay

One carbon elongation of the basic amino acid Lysine (by a −CH_2_ group) keeping other reactive groups (−COOH, −NH_2_ at one end and the −NH_2_ group at the other end) was carried out to examine the binding limit of the molecular chain length keeping the two ends untouched (Datta et al., 2001). As described (Materials and Methods above), the molecule was examined for angiogenic potential in cell culture assay with a cancer cell line A-549, expressing VEGFR to a very reasonable extent.

**Fig. 4.**
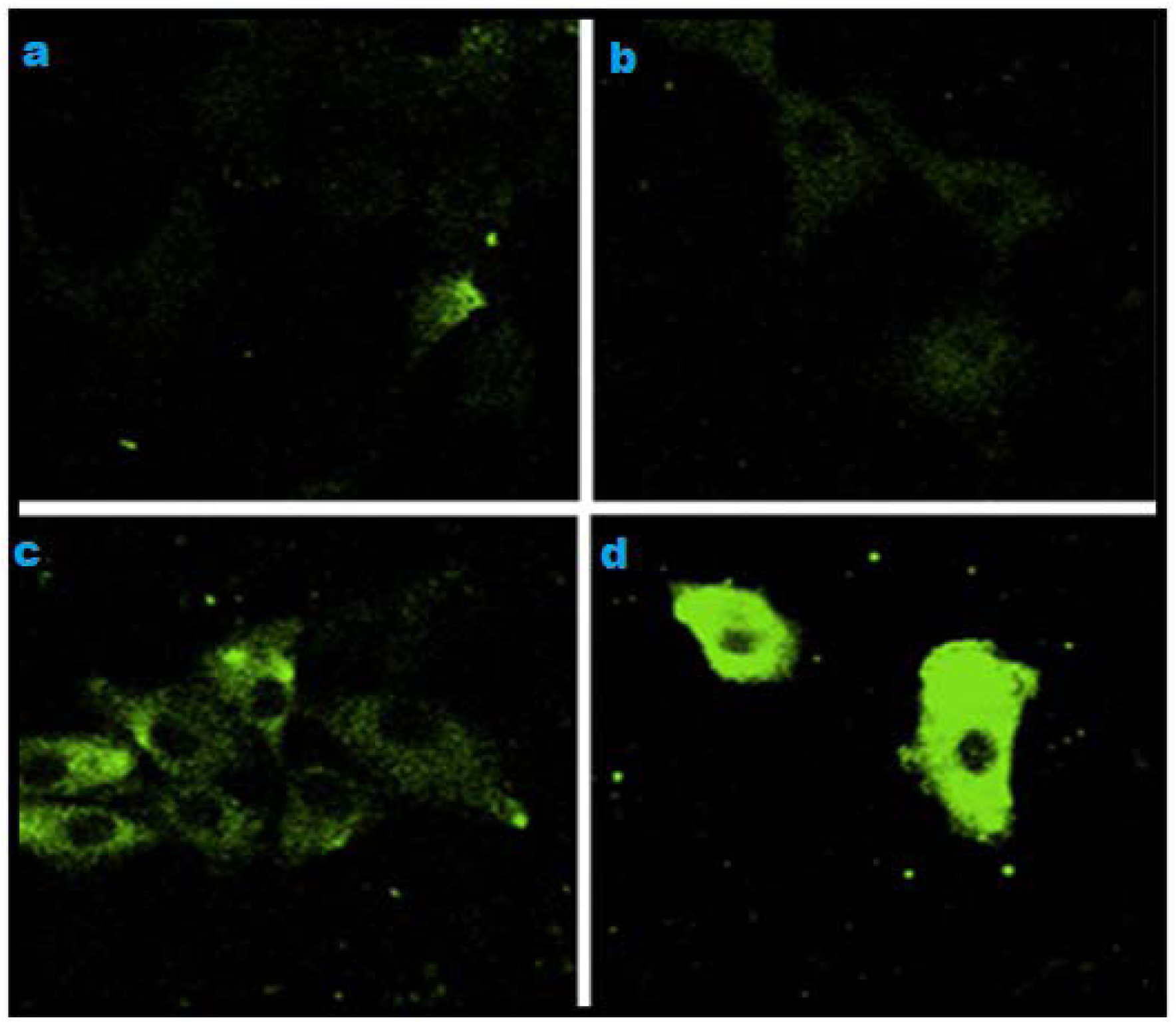
VEGF binding to VEGFR present on human lung epithelial cells untreated and treated with ELM: **a-** Growth medium only; **b-** Medium + ELM; **c-** Medium + VEGF; **d-** Medium + VEGF + ELM

Presence of VEGF only (Fig. 4c) in the medium gave a reasonable degree of fluorescence while the coupled presence of VEGF and the synthetic analogue enhanced the binding to a very remarkable extent (possibly exceeding by a few orders of magnitude) as seen in Fig. 4d. This definitely signifies a probable intervening binding role of the synthetic analogue molecule between VEGF and VEGFR (both unqualified).

## 3.3. In-silico docking studies of VEGF-A and VEGFR-1 binding in presence of ELM

Placement of the intervening molecule between VEGF-A and VEGFR-1 in two different orientations show distinct binding capabilities of the ELM molecule through its reactive –NH ^+^ groups at both ends of the molecule possibly “at a unique distance”, along with the -COOH group at the chiral carbon (Fig. 5).

**Fig. 5.**
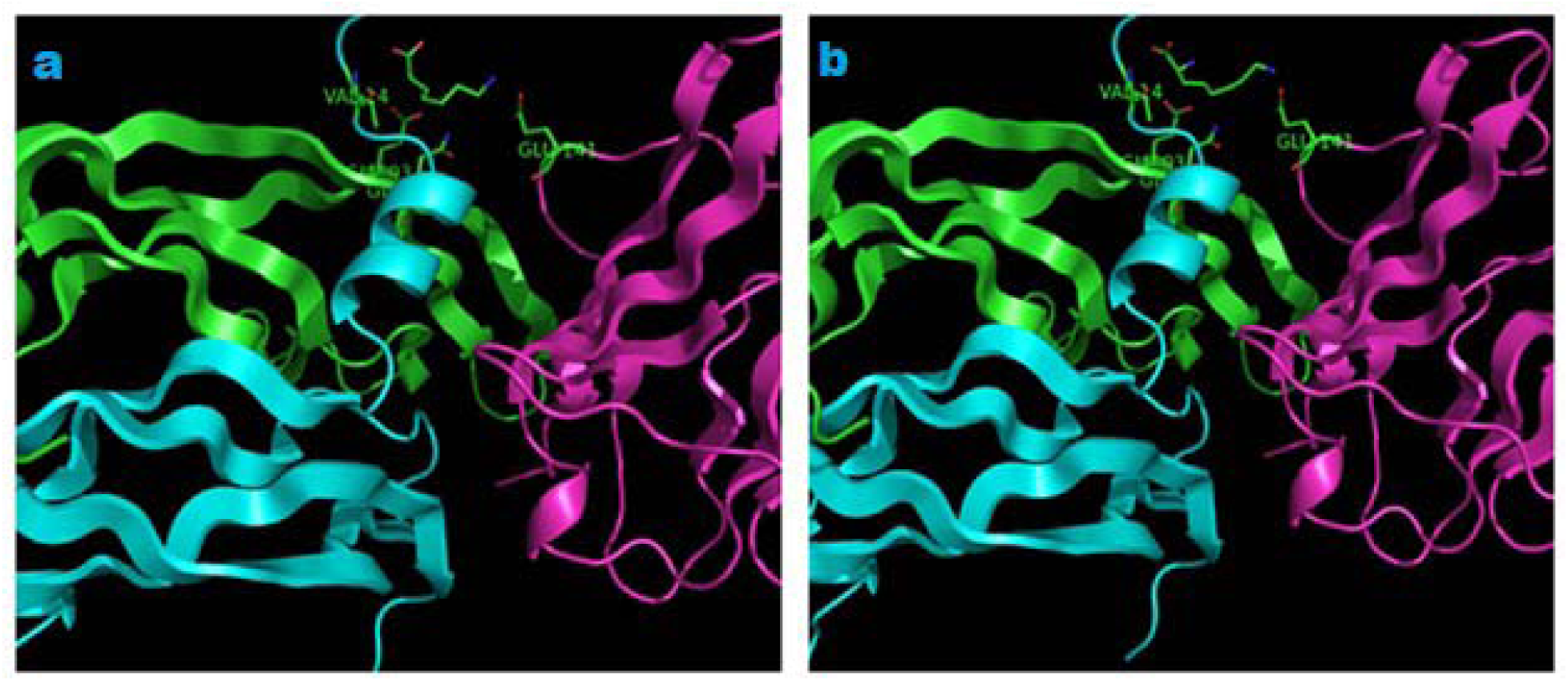
The molecular docking result showing the location of the foreign ELM molecule (on middle top) between VEGF-A (green and pink) and VEGFR-1(blue) with different orientations (**a** and **b**).

It will be of interest to examine the limiting length of the intervening binder molecule beyond which the biological response by way of induced angiogenesis is no more observed. In addition, it will be of interest to go back to the earlier description of preferential binding sites for added lysine moiety to both VEGFs and VEGFR(s). In both the peptides, despite a large distribution of negative charge loads on the respective molecule(s), both VEGFs and VEGFR(s) showed remarkable preferential binding site at unique and/or dispersed positions in most of the cases. Significance of such location(s) need further investigations.

## 4. Conclusions

Physiological angiogenesis being slow and nearly ineffective in all ischaemic conditions, needs augmentation for it to be functionally and clinically meaningful and as a significant mean of controlled reperfusion (Gupta et al., 2004). Since, it is mediated through VEGFs binding to their respective VEGFR(s), possible “anchorage capability of Lysine molecule” enhances the stability and associated biological response to a remarkable level (as discussed earlier).

Present study raises a crucial possibility of examination of other defined synthetic analogue(s) of the basic amino acid species as possible low molecular weight angiogen(s) in a family. Clinically, one or more of these (within a limiting chain length) may induce time-bound controlled angiogenic responses in ischaemic tissues and organs for faster therapeutic recovery of ischaemic conditions.

Base level binding stability of VEGFR-1 – VEGF-B versus the base level stability of VEGFR-1 – VEGF-A keeps one important question very open: Just being stable may not be the only answer to a comparable level of physiological action for a VEGF – VEGFR complex (angiogenic response). There may as well be some other controlling parameter(s) in the final phenotypic expression of the physiologic response. Also, it remains a standing question whether this Low-Mol-Wt natural and synthetic molecular species are mere stabilizers of the peptides’ bindings or do they individually modify some other molecular events leading to these augmented responses.

## Compliance with ethical standards

Biological studies had been conducted as per the institutional ethical committee guidelines and following all applicable regulatory norms.

